# Extensive plant use of exometabolites

**DOI:** 10.1101/2022.07.29.496484

**Authors:** Yuntao Hu, Peter F. Andeer, Qing Zheng, Suzanne M. Kosina, Kolby J. Jardine, Yezhang Ding, La Zhen Han, Yu Gao, Karsten Zengler, Benjamin P. Bowen, Jenny C. Mortimer, John P. Vogel, Trent R. Northen

**Author notes:** **Corresponding authors** Correspondence to Trent R. Northen.

## Abstract

Root exudation has been extensively studied due to its importance in soil carbon cycling and in supporting growth of soil microbes. However, the extent and dynamics of plant uptake of exogenous metabolites is poorly understood. To gain new insights into these processes we used ^13^C-tracing to characterize plant uptake of exometabolites across a panel of diverse plant species (*Arabidopsis thaliana, Brachypodium distachyon, Lotus japonicus, Panicum virgatum*, and *Kalanchoe fedtschenkoi*) grown in sterile hydroponic cultures. The uptake of exometabolites accounted for 23% of the overall *B. distachyon* carbon budget, and we identified 33 metabolites that were taken up by plants. Counterintuitively, many metabolites had higher uptake rates during the day vs. night. Thirteen of the metabolites from root exudates were found to promote root growth in *A. thaliana*, including hydroxybenzoate, threonate, *N*-acetyl-glucosamine, and uracil. Together these results indicate that the root uptake of organics can account for a significant portion of the plant carbon budget and that exogenous small molecules used by plants alter root growth with implications for plant nutrition, organic farming, soil nutrient cycling, and rhizosphere community dynamics.

## Main

Plant root exudation has attracted considerable interest given its contribution to soil carbon cycling, inorganic nutrient solubilization, and structuring rhizosphere microbial communities^1^. However, in principle these transport processes can be reversed, enabling roots to capture exogenous organic molecules. Indeed, there is evidence that plants can also uptake organic metabolites^2^ and that there are diurnal patterns of total organic carbon abundance in the rhizosphere^3^. Most work has focused on plant use of amino acids and peptides, which can serve as exclusive nitrogen sources^2,4-6^, though limited work has shown plants can use a larger range of metabolites^5,7^. This use of nitrogen-containing compounds is known to be economical for plants^8^ and can be especially important in ecosystems where ammonium and nitrate are limited^2,9^. Numerous membrane transporters associated with plant import of amino acids have been characterized, including LHT1, AAP1, and ProT2^10,11^. While plants can clearly uptake exogenous organic compounds, the extent and diversity of metabolites used, their dynamics and importance to plant carbon cycling, and their phenotypic effects remain poorly understood.

In this study we added ^13^C-labeled root extracts to hydroponic cultures of *B. distachyon* grown in sterile hydroponic medium using fabricated ecosystems (EcoFABs^12^) and measured significant ^13^CO_2_ respiration rates originating from these root extracts. A panel of diverse plants (*A. thaliana, B. distachyon, L. japonicus, P. virgatum*, and *K. fedtschenkoi*) was used to quantify tracer incorporation into plant tissues and the uptake of individual metabolites was analyzed. Furthermore, the dynamics of exogenous metabolites were investigated over the diurnal cycle in all plants. *A. thaliana* was used to characterize the nutritional and physiological effects of 26 of the most common metabolites in exudates.

### Plant uptake of exogenous metabolites can be a significant contribution to overall carbon flux

Soils contain a vast diversity of organic compounds^13^ many of which are thought to originate from plant roots^14^. Therefore, to investigate plant uptake and use of diverse exometabolites, we extracted root metabolites from the ^13^C-labeled grass *S. bicolor* (^13^C atom% > 92%), and ‘fed’ them to the small model grass *B. distachyon* grown in sterile EcoFABs while monitoring the release of ^13^CO_2_ (Fig. 1a, Fig. S1). We initially focused our analysis on the ^13^CO_2_ fluxes at “night” (dark period, 8h) because, during the “day” (light period, 16h), we found that the respired ^13^CO_2_ is partially re-assimilated by the leaves. At night, when photoassimilation processes ceased, we determined that ^13^CO_2_ was respired by the sterile plants at a rate of 0.7 *µ*mol ^13^CO_2_ g^-1^ DW h^-1^ (Fig. 1b), indicating some direct plant use of the exogenous metabolites at night as respiratory substrates, consistent with previous reports^15^.

**Figure 1.**
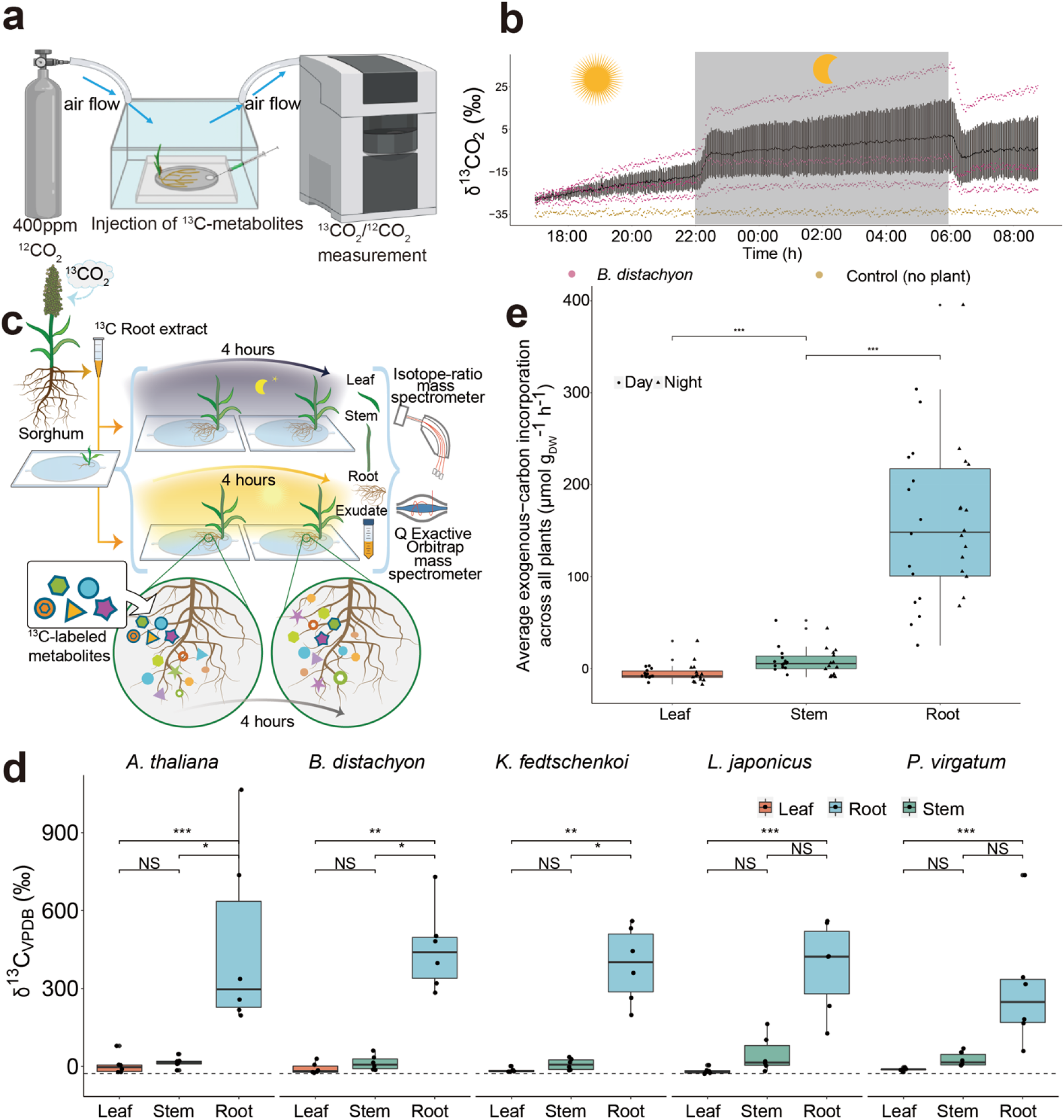
Isotopic tracing of respiration and plant use of ^13^C-labeled root extracts. **a**, Experimental setup used to introduce ^13^C-labeled exogenous metabolites and measure released ^13^CO_2._ **b**, Diurnal increase of δ^13^CO_2_ after injection of ^13^C root metabolites into the hydroponic growth medium. Black dots are the average δ^13^CO_2_ of EcoFABs with *B. distachyon* (n=3) with error bars (± 1 SE). Pink and brown dots are the raw measurement of δ^13^CO_2_ of EcoFABs with *B. distachyon* and empty EcoFABs (n=1). **c**, Experiment setup for the diurnal ^13^C isotope tracing assay. **d**, ^13^C enrichment in different plant tissues across all wild-type plants (n=6) (significances between roots, stems, and leaves were determined via Kruskal–Wallis test followed by Dunn’s test with Benjamini-Hochberg correction, NS. not significant, \**p*<0.05, \*\**p*<0.01, \*\*\**p*<0.001). The natural abundance (−27‰) of the ^13^C atom percentage in plants is indicated with a dashed line. **e**, Comparison of carbon incorporated into different plant tissues between day and night (n=15). See Methods for calculations. Significance between tissues was determined via Kruskal–Wallis test followed by Dunn’s test with Benjamini-Hochberg correction (\*\*\**p*<0.001).

To explore the range and extent of exogenous metabolite use across the diurnal cycle, we selected five divergent plant species (*A. thaliana, B. distachyon, L. japonicus, P. virgatum*, and *K. fedtschenkoi*), including representatives of monocots and eudicots, representing all three types of photosynthesis (C3, C4, and CAM) (Table S1). Plants were cultivated in sterile EcoFABs (Fig. 1c, Fig. S2) for three weeks, at which point, the same amount of ^13^C-labeled extract (equal to 20% of total carbon in the growth medium) of *S. bicolor* was added to each of the plants. The resulting dissolved organic carbon concentration of the plant growth medium ranged from 51 to 136 ppm (Table S3), comparable with the dissolved organic carbon in organic rich soils^16^. Nighttime experiments were started after 4-hour of darkness to ensure plants have started using the stored starch from photosynthesis^17^. The medium was then sampled following 1 and 4 hours of tracer addition. Following incubation with the tracer, the plants were removed and roots were washed 3 times with ionic buffer solution (50 mM CaCl_2_) to remove adsorbed metabolites from root surface. The spent medium samples from the EcoFABs were plated to confirm sterility and the metabolites were analyzed using liquid chromatography coupled tandem mass spectrometry (LC-MS/MS). Tissues were harvested, divided (roots, stems, and leaves) and analyzed for ^13^C using IRMS.

Significant increases in ^13^C-labeling were observed in both roots (δ^13^C, 399±40‰) and stems (δ^13^C, 22±7‰) across all five plants (Fig. 1d, Table S2) compared to the natural abundance - 27±2‰^18,19^. This can’t be solely explained by unspecific root sorption of exogenous metabolites as we observed respiratory production of ^13^CO_2_ emissions and an increase in high ^13^C% in stem tissue. Consistent with the short labeling period, we found much lower ^13^C-incorporation into the leaves and stems vs. the roots. We observed exogenous-carbon incorporation into plant biomass across the diurnal cycle and the five plants (Fig. 1e and Fig. S3) on average 160 *µ*mol C g^-1^ DW h^-1^. This high level of root carbon uptake from exogenous metabolites was consistent across the diverse species (Fig. 2a).

**Figure 2.**
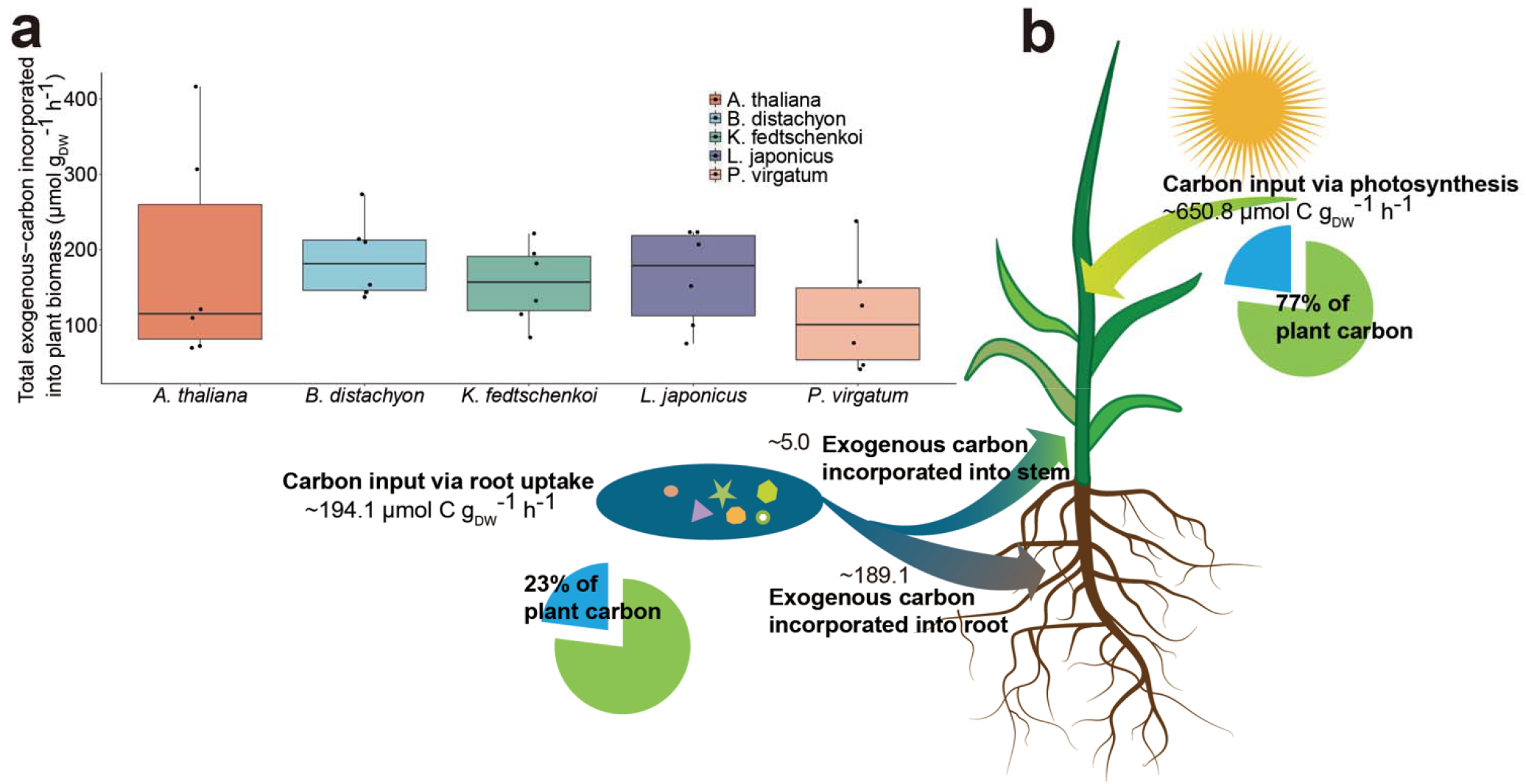
Contribution of exogenous organic carbon to the plant carbon budget. **a**, The exogenous carbon incorporated into plant tissues (stems and roots) across five divergent plant species during day and night (n=6). **b**, Schematic representation of the carbon flow through *B. distachyon* leaves and roots. Pie charts show the relative carbon flux via photosynthesis vs. exogenous-carbon use. Carbon input via photosynthesis was the average of photosynthesis rates measured by cavity ring-down spectroscopy and leaf-level gas exchange. Carbon input via root uptake was the sum of exogenous-carbon incorporated into plant tissues.

We determined the carbon budget for *B. distachyon* based on the tissue labeling results in combination with the net carbon fixation rate (photosynthetic – respiration rate). Here the photosynthetic rate was determined using two approaches, cavity ring-down spectroscopy and leaf-level gas exchange. The photosynthetic rates were determined to be 587 and 716 *µ*mol CO_2_ g^-1^ DW h^-1^, respectively. From this we found that 23% of *B. distachyon* carbon was coming from exometabolites, most of which was detected in the roots (Fig. 2b). Specifically, exogenous metabolite use amounted to 194 *µ*mol C g^-1^ DW h^-1^. Of this, 189 *µ*mol C g^-1^ DW h^-1^ was partitioned into root biomass, 5 *µ*mol C g^-1^ DW h^-1^ into stems (see Methods) and the rest was respired. There are few studies investigating the extent of plant use of exogenous organics that we can use for benchmarking these results: A ^14^C-tracing study found that rice assimilated 0.6 % of its total carbon from soybean manure^20^. Another study used ^13^C-tracing and found that 9% of the rice’s total carbon can be obtained from cattle manure, 40-70% of which was assimilated in the roots^21^. Hence, while we observe a similarly high partitioning into the roots, we detect a much higher contribution of exogenous carbon to the plant carbon budget (23% vs. 0.6-9%), which we attribute to multiple factors. First, the lack of microbial competition, transport, sorptive processes, etc. found in soils^2^. Second, the shorter incubation periods and low photosynthesis rate of young plants in our study. Finally, other studies would have primarily large, complex polymers as their extracellular component, whereas our study uses small molecules enabling direct plant uptake. Given these factors we expect that our values represent upper limits on plant use of exometabolites.

### Plants uptake diverse exogenous metabolites

To gain insights into which metabolites are being used by different plants we applied exometabolomics and ^13^C isotope tracing analysis^22,23^ (See Methods). Medium samples collected after 1 h and 4 h following the addition of the tracer were analyzed using LC-MS/MS-based metabolomics (Table S4). Ninety-eight metabolites were identified in the growth medium. Next, we determined the uptake of the exogenous metabolites based on the decrease of abundance of individual ^13^C-labeled tracer metabolite at 4 h vs. 1 h. We identified 33 metabolites in the exudates and were evaluated for the depletion of their ^13^C-labeled tracers. These ^13^C-labeled metabolites were all significantly depleted in the presence of the plants, including organic acids, nucleosides, in addition to the expected amino acids and sugars^2,5,24^. This is despite the fact that the inorganic nitrogen in the medium was sufficient to support plant growth (Table S5).

To examine the relative contributions of active vs. passive transport^25^ we further compared the abundance of root cell intracellular metabolite to that of root metabolite uptake (Fig. S4). For a passive process, metabolite uptake should be proportional to root tissue concentrations. However, we observed low r^2^-values suggesting active transport processes for the majority of metabolites, consistent with a previous study^5^.

### Plant use of exogenous metabolites varies across species and the diurnal cycle

Given indications of active transport processes (Fig. S4) and observed differences in ^13^CO_2_ production during the day vs. night, we compared the metabolomic data for significant diurnal cycle variation in plant use of individual metabolites. We observed clear differences in metabolite uptake across the five plants (Fig. 3a) which may be related to known differences in root exudate composition across species^26^. This could be partially caused by the different root surface as the apical zones have been shown to be the preferred metabolite exchange sites^27^. All plants showed a strong diurnal pattern of root metabolite fluxes (Fig. 3b and 3c). Amino acids formed a cluster of metabolites that were predominantly used during the day by all plants and metabolite fluxes also roughly clustered plants by species (Fig. 3b, 3c and S5). More detailed analysis revealed that *L. japonicus* (Fig. 3b) had a higher uptake for a subset of exogenous metabolites including amino acids and uridine during the day and that *K. fedtschenkoi* (Fig. 3c) had the highest uptake of asparagine and beta-alanine at night consistent with it being the only plant included in this study using crassulacean acid metabolism, where carbon is fixed at night^28^. We expected higher uptake of metabolites during the night when plants are not generating photosynthates. Surprisingly, many metabolites showed higher uptake during the day vs. the night (Fig. 3c and 3d). We speculate that the higher use of organics during the day may reflect a plant strategy to maintain C/N balance when photosynthesis is active^29,30^. Since non-N-containing metabolites may share the same transporters (e.g. GLUT transporters for both amino sugars and glucose)^31^, they may be indirectly used. Hence, the observed reduced exometabolite uptake at night may reflect the reduced nitrogen demand.

**Figure 3.**
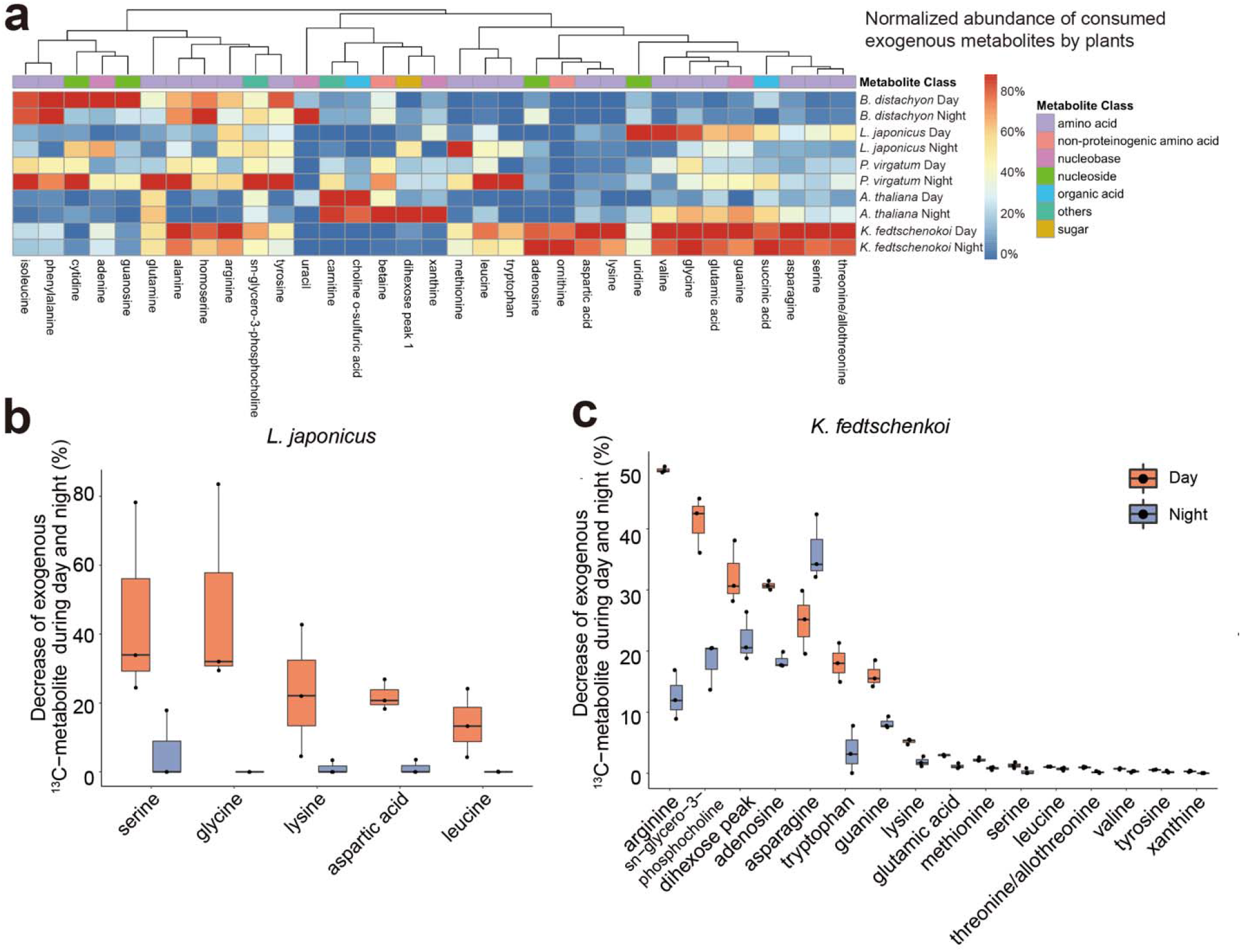
Dynamic root uptake of individual exogenous metabolites. **a**, Heatmap of normalized abundance (%) of consumed exogenous metabolites by roots across five plants during the day vs. night. The average peak height of metabolite was normalized by the maximum peak height across plants and diurnal cycle. **b**, Diurnal difference (*p*<0.05) of the decrease of individual exogenous ^13^C-labeled metabolites (%) in *L. japonicus* during the day (in orange) vs. night (in blue) (n=3, significance between day and night was determined via Kruskal–Wallis test). **c**, Diurnal difference (*p*<0.05) of the decrease of individual exogenous ^13^C-labeled metabolites (%) in *K. fedtschenkoi* during the day (in orange) vs. night (in blue) (n=3, significance between day and night was determined via Kruskal– Wallis test).

### Exogenous metabolites used by plants promote root growth

To explore the possible benefits of exogenous metabolites on plant phenotype, we selected a diverse panel of 24 metabolites that were found to be utilized by the plants (Fig. S6 and S7). These include several common amino acids (such as alanine, glycine, and arginine), which are well-known as alternative plant organic nitrogen sources^8^. We also included metabolites (thymine, threonic acid, ferulic acid, and hydroxybenzoic acid) that to our knowledge have not been shown to alter root growth. *A. thaliana* was grown on agar plates containing 0.5 mM of these individual metabolites in half-strength Murashige & Skoog medium with or without 0.5 mM NH_4_NO_3_ (to control for nitrogen-based effects). Under nitrogen deprivation conditions, we found that alanine and other nitrogen-containing metabolites, such as amino sugars (*N*-acetyl-glucosamine) and nucleobases (e.g., thymine, uracil), resulted in up to 50% longer roots (Fig. 4a and S6). These observations are consistent with previous studies showing that plants demonstrated the ability to uptake a variety of N-containing metabolites, such as amine, quaternary ammonium, and nucleobases^5,24^. Interestingly, root growth promotion was also observed when NH_4_NO_3_ is provided (Fig. S7). We attribute this to the direct bioavailability of the metabolites vs. nitrate, which must first be converted to ammonia and then glutamate^32^, making them efficient nitrogen sources.

**Figure 4.**
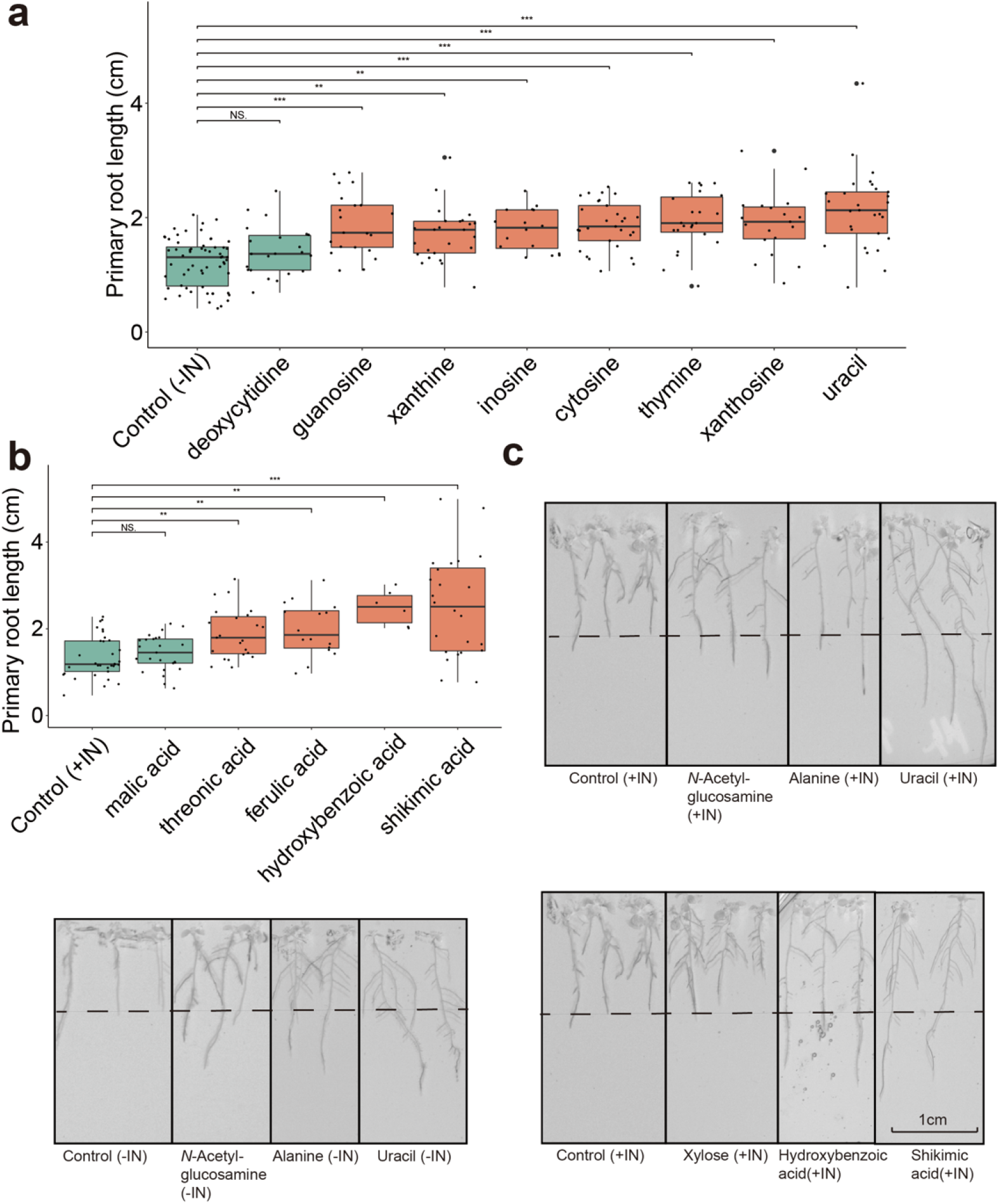
Growth promoting effects of exogenous metabolites used by plants. **a**, *A. thaliana* primary root length after 10 days of growth on plates containing 0.5 mM of metabolites and half-strength Murashige & Skoog medium lacking NH_4_NO_3_ (plants n=16 to 60, exact n number is in Table S8). Significance compared to control samples without metabolites was determined via Kruskal–Wallis test followed by Dunn’s test with Benjamini-Hochberg correction (NS. not significant, \*\**p*<0.01, \*\*\**p*<0.001). **b**, *A. thaliana* primary root length after 10 days of growth on plates containing 0.5 mM of metabolites and half-strength Murashige & Skoog medium with 0.5 mM NH_4_NO_3_ (plants n=6 to 30, exact n number is in Table S8). Significance compared to control samples was determined via Kruskal– Wallis test followed by Dunn’s test with Benjamini-Hochberg correction (NS. not significant, \*\**p*<0.01, \*\*\**p*<0.001). **c**, Selected pictures of the *A. thaliana* root under addition of selected metabolites (dotted lines represent a reference of the length of the primary root of the control group). First column: without inorganic nitrogen (-IN) NH_4_NO_3_; second columns: with 0.5 mM NH_4_NO_3_ (+IN).

Excitingly, even in the presence of excess ammonium nitrate, we observed root growth promotion by several metabolites that (to our knowledge) have not yet been observed to promote root growth. For example, the organic acids hydroxybenzoic acid and threonic acid both significantly promoted root growth and xylose increased the length of lateral roots (Fig. 4b and 4c). We also found that uracil-treated seedlings had nearly 2-fold longer primary roots (Fig. 4a). We next examined if mixtures of these metabolites would promote growth under non-sterile conditions. This was done by growing *B. distachyon* plants in pots filled with calcine clay treated with 0.5mM organic acids, 0.5-1mM nucleobases as compared to 1mM KNO_3_. It was found that plants supplemented with the organic mixtures had approximately twice the root biomass (Fig. S8). Furthermore, since nucleosides were found to promote plant growth and RNA is known to be an abundant nitrogen source in soils^33^ we examined if *A. thaliana* can use exogenous RNA as a nitrogen source. While we did not find RNase activity in the growth medium of *A. thaliana*, plants had significantly longer roots vs. control when treated with a mixture of RNA (containing 0.5 mM N) and RNase (Fig. S9), suggesting that plants can indeed use depolymerized RNA, presumably as a nitrogen source.

While soil organics have long been associated with fertility, this is typically attributed to a range of benefits, from supporting microbial diversity to retaining nutrients, that do not include being used by plants. While it is likely that the contribution of organics to the plant carbon budget measured here represents upper limits, we speculate that the direct use of organics could be particularly beneficial for young plants. This may be especially important when photosynthesis is limited, for example by shading from more mature plants. These findings suggest a direct connection between fertility and the concentration of organics in the environment, also supported by limited field studies^20,21^.

Overall, we observed that diverse exogenous metabolites we identified were taken up by a diverse panel of plants grown in sterile hydroponic conditions and that the captured exogenous carbon accounted for 23% of the overall plant carbon budget. This uptake varied between plant species and across the diurnal cycle. Many of the metabolites used by plants were found to promote root growth. Together these findings highlight the potential contribution of plant organic use to plant growth and nutrition with important implications to carbon cycling, microbial interactions, and plant nutrition.

## Methods

### Preparation of ^13^C-labeled metabolite tracers (uniformly ^13^C-labeled root metabolites)

Sorghum seeds, kindly provided by Professor Jeff Dahlberg (UCANR), were germinated on soil under controlled conditions (30 °C, 30% humidity, 150 *µ*mol m^-2^ s^-1^, 14 h light/day) for 17 days. Seedlings with four leaves were then transferred into a hydroponic container with Hoagland’s nutrient solution (pH 6.0) and grown in a self-constructed growth chamber^34^ with a controlled environment (air with 600 ppm ^13^CO_2_, 28 °C, 70% humidity, 155 *µ*mol m^-2^ s^-1^, 18 h light/day) for ^13^C enrichment. After an additional 29 days of growth, plants were harvested and directly frozen in liquid nitrogen. The stem tissues were collected for ^13^C incorporation measurements, and the plants were then stored at -80 °C for later analysis. The ^13^C incorporation measurements were conducted by analyzing the oligosaccharides released from digestion of the stem samples with endo-1,4-□-xylanase (Megazyme, rumen microorganism, GH11) by matrix-assisted laser desorption/ionization-time of flight (MALDI-TOF) mass spectrometry (UltrafleXtreme instrument, Bruker) as previously described^35^. The oligosaccharide, XAXX^36^, was used to determine the ^13^C incorporation, which is 92% for this study. ^13^C-labeled root metabolites were prepared by extracting lyophilized sorghum roots with methanol^12^. The methanolic extracts were dried and stored at -80 °C.

### Plant ^13^CO_2_ respiration experiment

Analysis of the stable carbon isotope composition of CO_2_ (δ^13^CO_2_) and absolute ^13^CO_2_ mixing ratios (ppm) was performed in real-time using a cavity ringdown spectrometer (CRDS, G2131-i, Picarro Inc., Santa Clara, CA, USA). The CRDS was factory calibrated for high precision δ^13^CO_2_ (< 0.1 ‰) at 400 ppm CO_2_. *B. distachyon* 21-3 was obtained from the Joint Genome Institute *Brachypodium distachyon* collection. Plant seeds were sterilized in 6% v/v sodium hypochlorite and 70% v/v ethanol for 30 seconds and 5 mins, respectively. All seeds were then washed five times in sterile water and kept in the dark at 4 °C for 5 days prior to the inoculation. Seedlings were germinated on agar plates (half-strength Murashige & Skoog salts^34^ with 1.5% agar) for 2-5 days and transferred to EcoFABs after germination. EcoFABs made of polycarbonate (EcoFAB 2.0) were autoclaved, assembled, and planted with *B. distachyon* seedlings (Fig. S1) with liquid half-strength Murashige & Skoog medium^37^. Murashige & Skoog medium was prepared by dissolving Murashige & Skoog inorganic salts without the addition of micro-organic nutrients. The assembled EcoFABs consisted of three polycarbonate pieces, a glass slide, and a silicone gasket to generate two chambers, one for the plant shoots and the second for roots, separated by a port for the seedling. The top polycarbonate chamber consisted of ∼ 50 mm of height and 253 cm^3^ of volume for plant shoot growth and the root chamber provided 2 mm of depth and ∼ 12 mL volume for plant root growth. Individual EcoFABs (empty with buffer, and with buffer and plant) were placed in a growth chamber (E36HO, Percival Scientific) with constant air temperature (25 °C) and daytime (6:00-22:00) photosynthetically active radiation flux of 10 to 15 *µ*mol m^-2^ s^-1^, at the top of the EcoFAB. 25 mL/min of the EcoFAB headspace atmosphere was continuously withdrawn from the headspace output port and directed to the inlet of the CRDS for analysis using 1/8” O.D. Teflon (PTFE) tubing. In order to eliminate potential spectral interferences of the CO_2_ isotopic measurements, and to prevent CRDS damage from condensation, water vapor was first removed just prior to entering the CRDS inlet by first passing through a water trap (Chemical scrub tube assay, part 9960-093, Licor Inc.) packed with blue indicating Dririte (10-20 mesh, Licor Inc.). Due to the higher volume of sample path caused by the water vapor trap, a CRDS response delay of ∼5 min was determined from when changes in EcoFAB headspace δ^13^CO_2_ and ^13^CO_2_ were recorded by the CRDS. Air leaving the EcoFAB for isotopic analysis of CO_2_ was continuously replaced with 400 ppm certified CO_2_ standard in the air (Praxair, Inc.). To accomplish this, 100 mL/min of the 400 ppm CO_2_ standard was continuously delivered to a 1/8” O.D. stainless steel tee (Swagelok) just upstream of the EcoFAB air inlet via 1/8” O.D. Teflon (PTFE) tubing with 25 mL/min entering the EcoFAB and the remaining 75 mL/min venting to the growth chamber. In this way, air flows into and out of the EcoFAB were in balance which ensured that the air pressure inside the EcoFAB remained identical to that of ambient air inside the growth chamber. Following 1 hour of CRDS measurements of an empty EcoFAB, the gas inlet and outlet tubing, and fittings were quickly switched to an EcoFAB with buffer (control) or EcoFAB with buffer and plant. Following 1 hour of CRDS measurements, 2 mL of the ^13^C-labeled substrate solution was injected into the EcoFAB liquid port/septa using a needle with syringe. The final tracer concentration in the growth medium is 16 ppm C, which is approximately 20% of the EcoFAB root chamber TOC concentration. Continuous CRDS measurements of the dynamic EcoFAB headspace ^12^CO_2_ and ^13^CO_2_ mixing ratios (ppm) and δ^13^CO_2_ (‰) continued for 5-6 days. Three replicate plant experiments were performed. The sterility of the growth medium was checked by spreading one drop of growth medium on half-strength LB agar plates to ensure no growing microbial colony developed. In contrast to EcoFABs with buffer and plant, control EcoFAB experiments with buffer and ^13^C tracers only (no plant) did not show detectable ^13^C enrichment of headspace CO_2_, which retained the δ^13^CO_2_ of the 400 ppm CO_2_ standard (∼ -34‰).

### Plant photosynthesis experiments

The photosynthesis rate of *B. distachyon* 21-3 (3-week seedling) was first measured using a portable photosynthesis system (Li6800 with light source 6800-02, Li-Cor Biosciences, USA) installed with a 6 cm^2^ leaf chamber aperture. *B. distachyon* 21-3 was germinated and cultivated in polycarbonate EcoFABs as described above. The leaf chamber was set to 25 °C, with 400 ppm reference CO_2_ at a flow rate of 131 *µ*mol s^-1^ through the sample chamber. The light was held at 150 *µ*mol m^-2^ s^-1^. Since the *B. distachyon* 21-3 cannot fill the 6 cm^2^ leaf chamber, the leaf area was estimated using Image J for comparison with literature values. The photosynthesis rate was then measured using the CRDS and the growth chamber mentioned above in the plant respiration experiment section, but with a 40 mL/min gas flow rate and under 150 *µ*mol m^-2^ s^-1^ illumination.

### Plant materials and growth conditions

*Arabidopsis thaliana* Col-0, *Lotus japonicus, Panicum virgatum*, and *Kalanchoe fedtschenkoi* were obtained from the Arabidopsis Biological Resource Center (OH, USA), Joint Genome Institute *Brachypodium distachyon* collection, National Plant Germplasm System (GRIN, MD, USA), and Succulentsdepot (https://www.succulentsdepot.com/), respectively. Plants were cultivated in triplicates in the PDMS EcoFABs, which are growth chambers consisting of 3D printed polydimethylsiloxane (SYLGARD, Dow) and microscope slides as described elsewhere^12^. The EcoFAB includes a plant reservoir for a single seedling and two ports for medium exchange under sterile conditions. The EcoFABs were prepared as previously reported and sanitized by rinsing with 70% ethanol and 100% ethanol^12,38^. Plant seeds, except for *K. fedtschenkoi*, were sterilized in 6% v/v sodium hypochlorite and 70% v/v ethanol for 30 seconds and 5 mins, respectively. All seeds were then washed five times in sterile water and kept in the dark at 4 °C for 5 days prior to the inoculation. Seedlings were germinated on agar plates (half-strength Murashige & Skoog salts^37^ with 1.5% agar) for 2-5 days and transferred to EcoFABs after germination. *K. fedtschenkoi* plantlets were propagated from leaf cuttings, where the original leaves were sterilized by wiping all the surfaces with 70% v/v ethanol for 2 mins. Plants with similar sizes were transferred into the EcoFABs for 1 week to allow the growth of roots. Plants were cultivated in liquid half-strength Murashige & Skoog medium^37^ in a growth chamber at 24 °C with 150 *µ*mol m^-2^ s^-1^ illumination (16 h light and 8 h dark). The growth medium was replenished every 2-3 days depending on the loss of medium in the EcoFABs due to evaporation and transpiration, and final concentration of ammonium and nitrate in the medium was measured according to a photometric approach by treatment with 2,6-dimethylphenol^39^ (Table S5). To determine how much tracers should be added, we cultivated an extra eight *B. distachyon* in the EcoFABs for 21 days and measured the total organic carbon (TOC) with Shimadzu TOC-L Analyzer and the average TOC in the growth medium is 87 ppm (Table S3).

### Root exogenous metabolite diurnal uptake measurement

After 21 days of cultivation, all plants were divided into day and night groups in triplicates. All the later experimental procedures were conducted in the dark for the night group and in the light for the day group. Tracer addition was conducted on the same day but at a different time for the two groups. Uniformly ^13^C-labeled root metabolites were redissolved in water and diluted to reach approximately 20% of the total organic carbon (16 ppm) in the plant growth medium on the 21st day. In the night group, 0.25 mL ^13^C-labeled tracer were injected into the growth medium to label the plant exogenous metabolite pool 4 h after the light was turned off, while in the day group ^13^C-labeled tracers were added 4 h after the plant entered the light cycle. The sterility of growth medium was checked by spreading ∼20 *µ*l of growth medium on the half-strength LB agar plates to ensure no growing microbial colony developed. These plates were inspected after three days and revealed no microbial growth. The EcoFABs were gently shaken for 15 s, allowing a homogeneous mix of the ^13^C-labeled tracers and exogenous metabolite pool. 1 h (t_1_) and 4 h (t_2_) after the injection of the tracers, 0.5 mL of the growth medium was sampled for LC-MS/MS analysis. Exogenous metabolite samples were immediately stored at -80 °C. The exogenous metabolites were further lyophilized for 48h, resuspended in 1 mL of methanol, vortexed for 30 s, and sonicated for 15 mins. Methanol extracts were centrifuged at 9000g for 3 mins to remove insoluble components and the supernatants were stored at -80 °C for LC-MS/MS analysis^12^.

### ^13^C enrichment in plant tissues

Lyophilized roots, shoots, and leaves were analyzed for ^13^C isotopes at the University of California, Davis Stable Isotope Facility (https://stableisotopefacility.ucdavis.edu/). This facility uses a PDZ Europa ANCA-GSL elemental analyzer coupled to a PDZ Europa 20-20 isotope ratio mass spectrometer (Sercon Ltd., Cheshire, UK), which has a long-term standard deviation is 0.2 per mil for ^13^C. Samples were firstly oxidized in a combustion reactor filled with Cr_2_O_3_ and silvered CuO at 1000 °C and then reduced in a reduction reactor packed with reduced copper at 650°C. The carrier gas (He) was dried by a water trap with Mg(ClO_4_)_2_ and P_2_O_5_. A Carbosieve GC column (65 °C, 65 mL/min) was used for the separation of N_2_ and CO_2_. At least four different laboratory stable isotope reference materials were included in the sample sequence. These materials have been calibrated against Standard Reference Materials (USGS-40, USGS-41, USGS-42, USGS-43, USGS-61, USGS-64, USGS-65, and IAEM-600). The provisional isotope ratios of all the samples are measured relative to the reference gas peaks from each individual sample. After correcting the values of the whole sample sequence using the known values of the laboratory reference materials, these provisional values were finalized by the stable isotope facility.

### Root metabolite extraction

Plants were removed from the EcoFABs, and the live roots were washed three times with 15 mL 50 mM CaCl_2_ to remove any residues attached to the root surface by dipping them in the solution for 10 s. All plant tissues were lyophilized for 48 h and weighed. Lyophilized roots were ground with a bead-beater for 5 s three times using 2 mm steel balls and the powder was extracted in 3 mL methanol, which was vortexed for 30 s and sonicated for 15 mins. Methanol extracts were centrifuged at 9000g for 3 mins to remove insoluble components and supernatants were stored at -80 °C until further LC-MS/MS analysis^12^.

### Metabolomics analysis

Methanol extracts of the root exogenous metabolite samples and root tissue samples were dried using a Speed Vac Vacuum Concentrator (Thermo) for 12 h. Dried samples were then resuspended in 150 *µ*L methanol containing internal standards (15 *µ*M U-^13^C, ^15^N amino acid mix, 5 *µ*g/mL ^13^C_2_, ^15^N_3_ cytosine (Sigma), 10 *µ*g/mL U-^13^C mannitol and trehalose (Omicron), 4 *µ*g/mL U-^15^N adenine, 3 *µ*g/mL U-^15^N hypoxanthine, 2 *µ*g/mL U-^13^C, ^15^N uracil, 5.5 *µ*g/mL U-^15^N inosine (CIL)) and filtered through 0.22 *µ*m centrifugal tubes (Nanosep MF, Pall Corporation, NY). Polar metabolites were chromatographically separated using hydrophilic interaction liquid chromatography using an InfinityLab Poroshell 120 HILIC-Z column (Agilent) on a 1290 UHPLC (Agilent) and analyzed on a Q Exactive Hybrid Quadrupole-Orbitrap Mass Spectrometer (Thermo Fisher Scientific, San Jose, CA, USA). Detailed LC and mass spectrometry conditions are provided in Table S4. Metabolite data were analyzed using Metatlas toolbox to obtain the extracted ion chromatograms and peak height^38^. Unlabeled metabolites were first identified as described in SI Table S4; briefly sample m/z values, MS/MS spectra, and retention times (RTs) were compared with an in-house library of authentic reference standards (Table S6). We then analyzed the samples containing labeled sorghum root extracts for the ^13^C-labeled isotopologues of these metabolites based on the RTs and theoretical m/z of the adduct detected in the unlabeled samples. The internal standards were used for correcting retention times across samples. We also used the internal standards to check for matrix effects (signal suppression from co-eluting interferences) as indicated by significant decreases in intensity between time-points within the retention times of the identified metabolites. Alanine (RT: 12.4-9 min) in the *P. virgatum* samples had a small (1.2-fold) but significant decreases. This is the same range as two identified metabolites, carnitine (RT: 12.6 min) and threonine (RT: 12.8 min) and while the decrease in these two tracers was larger (1.4 and 2.2, respectively) matrix effects may contribute to the observed decrease of these two metabolites by *P. virgatum*.

### Root exogenous metabolite flux and carbon incorporation calculations

Plant respiration of ^13^C-labeled tracers at night was calculated based on

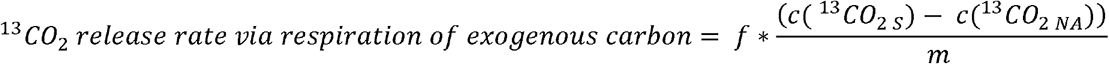

Where f is the flow rate of the air, ^13^CO_2 S_ is the ^13^CO_2_ concentration in the EcoFABs with plants, ^13^CO_2 NA_ is the natural abundance of ^13^CO_2_, and m is the biomass of plants.

Plant net photosynthesis rate during the day measured with portable photosynthesis system (Li6800, Li-Cor Biosciences, USA) was calculated based on

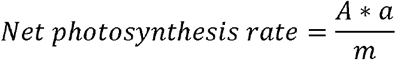

Where A is the net photosynthesis rate, a is the area of the leaf chamber aperture (6 cm^2^), and m is the biomass of plants.

Plant net photosynthesis rate during the day measured with CRDS (G2131-i, Picarro Inc., USA) was calculated based on

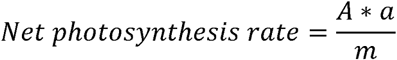

Where f is the flow rate of the air, ^12^CO_2 std_ is the ^12^CO_2_ concentration in the CO_2_ standard, ^12^CO_2 S_ is the ^12^CO_2_ concentration in the EcoFABs with plants, and m is the biomass of plants.

The amount of total carbon from exogenous metabolites incorporated into different plant tissues was calculated as follows:

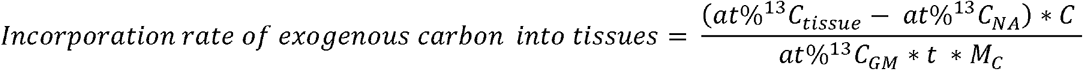

Where at%^13^C_tissue_ and at%^13^C_GM_ represent the ^13^C atom% in the plant tissues and growth medium. And at%^13^C_NA_ is the natural abundance of ^13^C atom% in plants^18^. C is the carbon content per gram of plant biomass. t is the incubation time (4 h) as described above, and M_C_ is the molar mass of carbon (12 g/mol). All the IRMS data and calculations are shown in Table S2. Root uptake of each metabolite was calculated by subtracting the peak height of exogenous ^13^C-metabolites of t_2_ from t_1_.

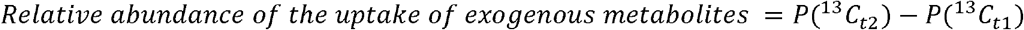

Where P(^13^C_t1_) and P(^13^C_t2_) represent the peak height of exogenous ^13^C-labeled metabolites. The termination times t_1_ and t_2_ were set to 1 h and 4 h according to experimental sampling times. All the calculation steps are included in Table S7.

### Effect of individual captured metabolites on plant growth under sterile condition

Based on the findings of plant uptake of metabolites contained in root exogenous metabolites, we selected metabolites to test their effect on the growth of *A. thaliana*. A total of 24 metabolites that were found to be significantly used by one or more plants were selected to span chemical diversity. First, we mixed 500 *µ*M of these metabolites with half-strength Murashige & Skoog medium lacking inorganic nitrogen to test the growth promotion effect of these metabolites under nitrogen deprivation conditions. In addition, we also mixed 500*µ*M of these metabolites with half-strength Murashige & Skoog medium supplemented with 500 *µ*M NH_4_NO_3_ to test the growth promotion effect of these metabolites when sufficient inorganic nitrogen is present. Controls with additional inorganic nitrogen added to half-strength Murashige & Skoog medium (1 mM and 5 mM NH_4_NO_3_) were used to test the growth promotion effect of additional inorganic nitrogen. The pH of all the medium were adjusted to 5.5. *A. thaliana* seeds were sterilized as described before and were germinated on these agar plates (15 replicates) in a growth chamber at 24 °C with 150 *µ*mol m^-2^ s^-1^ illumination (16 h light and 8 h dark). On the 11^th^ day after the seeds were exposed to light, the plants were imaged using a scanner and root hairs were visualized under a Leica M165FC microscope. Root images were analyzed using Image J with the smart root plugin.

### Effect of captured metabolites mixture on plant growth under non-sterile condition

According to the growth promotion effect of individual exogenous metabolites, we selected three organic acids and nucleobases to compare their growth promotion effects with nitrate on *B. distachyon* in non-sterile pot experiments. *B. distachyon* was washed as described above and germinated in 4-inch pots filled with porous ceramic (Profile Inc., Conover, NC, USA). Plants were watered every 4 days and supplemented with 150 mL Murashige & Skoog medium under 20 h / 4 h light cycle at 24°C. In addition, the controlled group was supplemented with 150 mL of 1 mM KNO_3_ every 4 days. The organic acids group and nucleobases group were supplemented with 0.5 mM organic acids mixture and 0.5 mM / 1 mM nucleobases mixture every 4 days. 0.5 mM organic acid mixture contains 0.17 mM of threonic acid (Sigma), 0.17 mM of hydroxybenzoic acid (Sigma) and 0.17 mM of shikimic acid (Sigma) (0.5 mM organic acid in total). 0.5 mM nucleobase mixture contains 0.17 mM of uracil (Sigma), 0.17 mM of thymine (Sigma) and 0.17 mM of cytosine (Sigma) (0.5 mM nucleobase in total). 1 mM nucleobase mixture contains 0.33 mM of uracil, 0.33 mM of thymine and 0.33 mM of cytosine (1 mM nucleobase in total). Plants were harvested after 2 weeks and the root biomass was recorded.

### Plant utilization of exogenous RNA

Based on the findings of plant uptake nucleotides, we tested the growth promotion effect of exogenous RNA and RNA plus RNase on *A. thaliana*. We mixed exogenous RNA consisting of 500 *µ*M of nitrogen, 70 *µ*U/mL of Ribonuclease A (EC 3.1.27.5) from bovine pancreas, and half-strength Murashige & Skoog medium lacking inorganic nitrogen as the growth medium. The pH of the medium was adjusted to 5.5. *A. thaliana* seeds were sterilized as described before and were germinated on these agar plates (15 replicates) in a growth chamber at 24 °C with 150 *µ*mol m^-2^ s^-1^ illumination (16 h light and 8 h dark). 11 days after the seeds were exposed to light, the plants were imaged using a scanner and root hairs were visualized under a Leica M165FC microscope. Root images were analyzed using Image J with the smart root plugin.

### Statistical analysis

All statistics and calculations were conducted with R (version 3.6.2). Data were tested for homogeneity of variance and transformed if necessary. The difference between groups was tested by Kruskal–Wallis tests followed by Dunn’s test with Benjamini-Hochberg correction (Fig. 1d, 1e, 4a, 4b, S3, S6, S7, and S8). Significance was indicated by NS. not significant, \**p*<0.05, \*\**p*<0.01, \*\*\**p*<0.001. Additionally, non-metric multidimensional scaling (NMDS) analysis was performed on the root exogenous metabolites uptake of different plant species. Hierarchical clustering heatmap (Fig. 3a) was analyzed based on Euclidean distance using R ‘pheatmap’ package 1.0.12.

## Supporting information

All SI figures

## Data availability

All the raw LC-MS/MS data included in this study have been deposited to the MassIVE.quant platform (https://massive.ucsd.edu/ProteoSAFe/index.jsp) from the University of California, San Diego, under ID MSV000088644.

## Code availability

The source code of the Metatlas toolbox used to obtain the extracted ion chromatograms and peak height in this study is openly available from the GitHub repository accessible at https://github.com/biorack/metatlas.

## Author Contributions

Y.H. and T.R.N. designed the project. Y.H. and Q.Z. conducted the isotope tracing experiment. K.J.J. conducted the respiration experiment. Y.G. and J.M. harvested and characterized ^13^C-labeled Sorghum roots. J.P.V. provided and cultivated model plants. Y.H., S.M.K., and B.P.B. analyzed the LCMS data and IRMS data. Y.H., P.F.A., B.P.B., and T.R.N. analyzed all the data and created figures. Y.H., P.F.A., S.M.K., Y.Z., L.Z.H., K.Z., T.R.N. wrote the manuscript with significant input from all other authors.

