## Supplementary material for "Extensive plant use of exometabolites": All SI figures

### Supplementary figures:

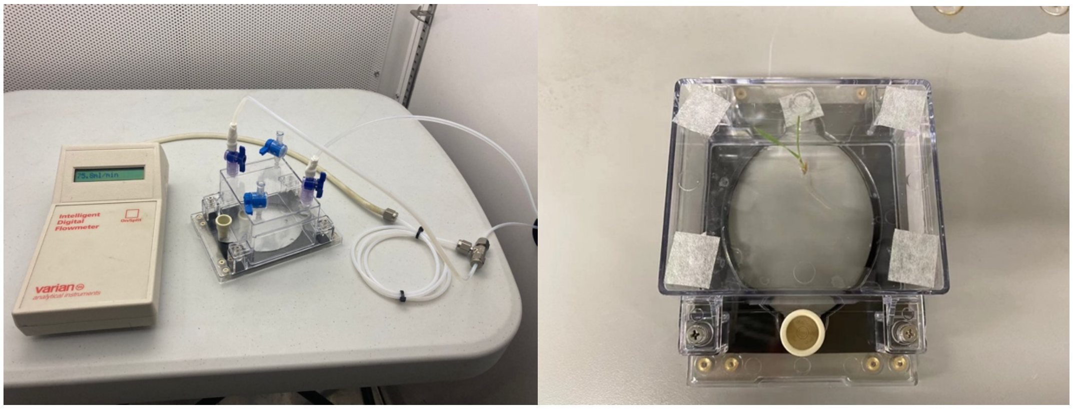

**Figure S1 Experimental setup for the measurement of stable carbon isotope composition of plants releasing $CO_2$ in the EcoFABs. a**, Flow meter between the EcoFAB and the gas sampling tube. **b**, EcoFAB 2.0 with *B. distachyon* inside.

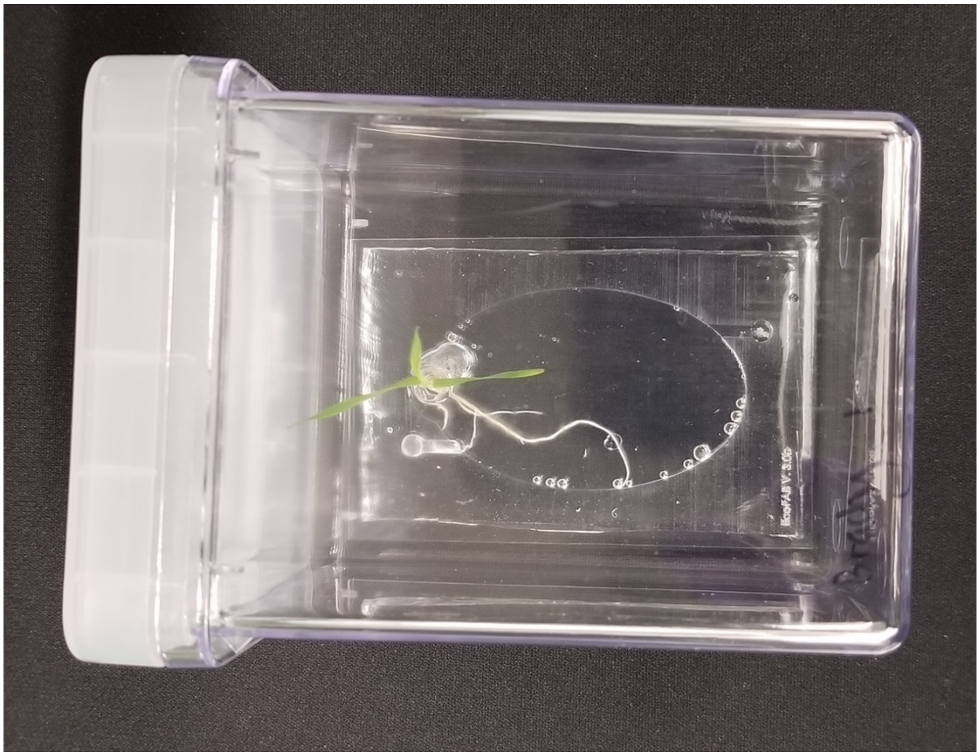

**Figure S2 Example of *B. distachyon* grown in PDMS EcoFABs**

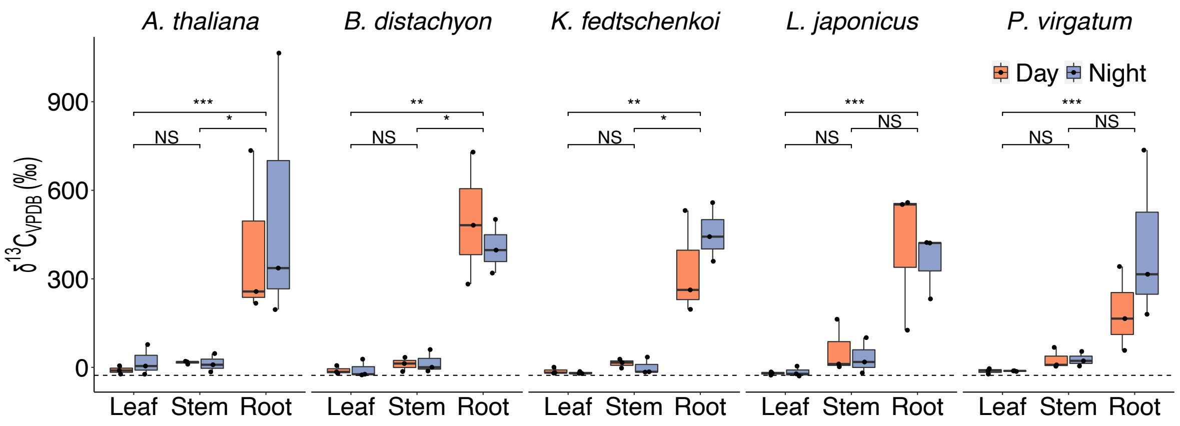


**Figure S3 $^{13}$C atom percentage in different plant tissues across all wild-type plants (n=3).** The natural abundance range of the $^{13}$C atom percentage in plants is indicated with dashed lines. The data shown in this figure is identical to figure 1d, but grouped by tissues and day vs. night. Significance between tissues was determined via Kruskal–Wallis test followed by Dunn's test with Benjamini-Hochberg correction, NS. not significant, *$p$<0.05, **$p$<0.01, ***$p$<0.001.

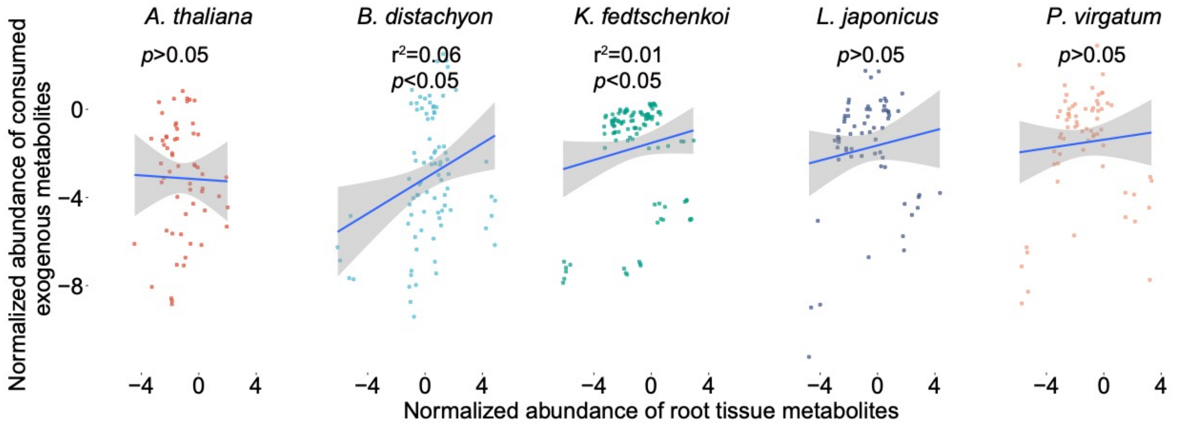


**Figure S4** Spearman correlation analysis of normalized abundance (peak height of individual metabolite is normalized to its corresponding internal standard) of consumed $^{13}$C-labeled exogenous metabolites by plants and normalized abundance (peak height of individual metabolite is normalized to its corresponding internal standard) of root intracellular metabolite across all wild-type plants (n=6). Only metabolites with corresponding internal standards are included, which are adenine, alanine, arginine, asparagine, aspartic acid, glutamic acid, glutamine, glycine, isoleucine, leucine, lysine, methionine, phenylalanine, serine, threonine/allothreonine, tryptophan, tyrosine, uracil, and valine.

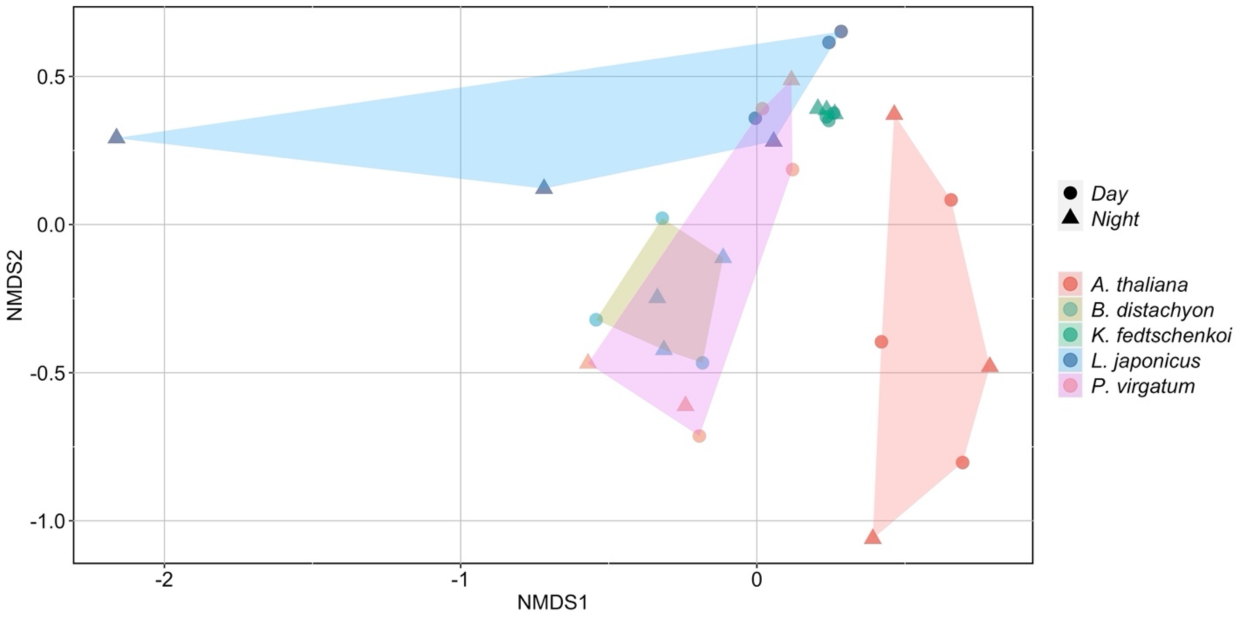


**Figure S5 NMDS analysis of consumed $^{13}$C-labeled exogenous metabolites by roots across five plant species and the diurnal cycle.**

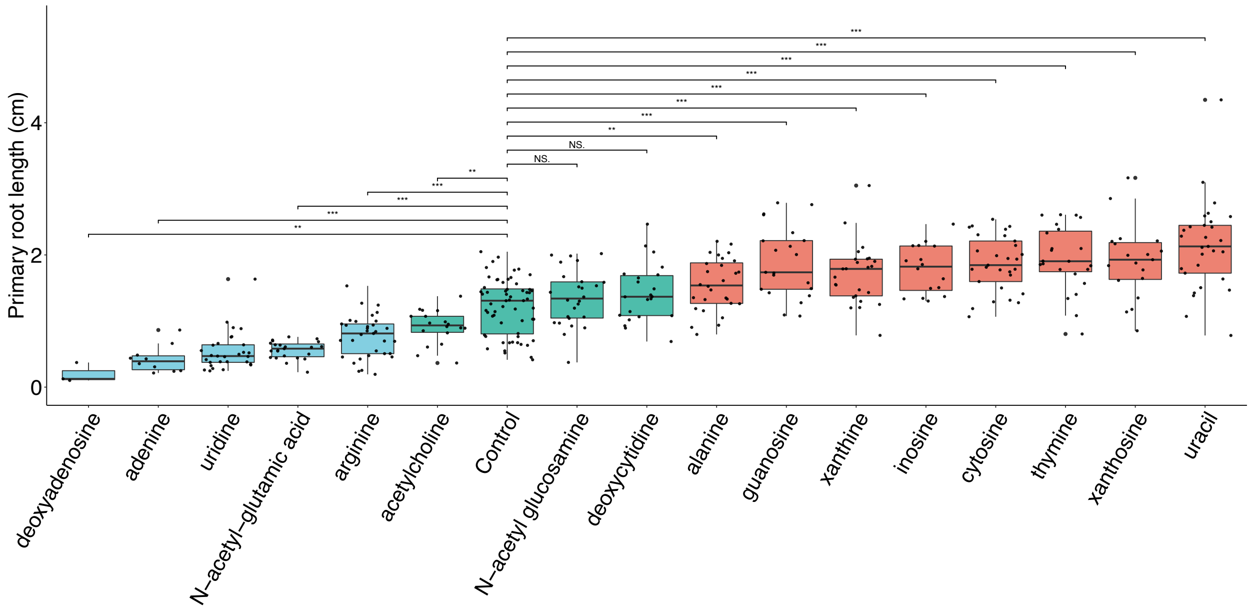


**Figure S6 *A. thaliana* primary root length under addition of 0.5 mM selected metabolites under nitrogen deprivation conditions (n=19 to 31).** Significance was determined via Kruskal–Wallis test followed by Dunn's test with Benjamini-Hochberg correction (NS. not significant, **p*<0.05, ***p*<0.01, ****p*<0.001).

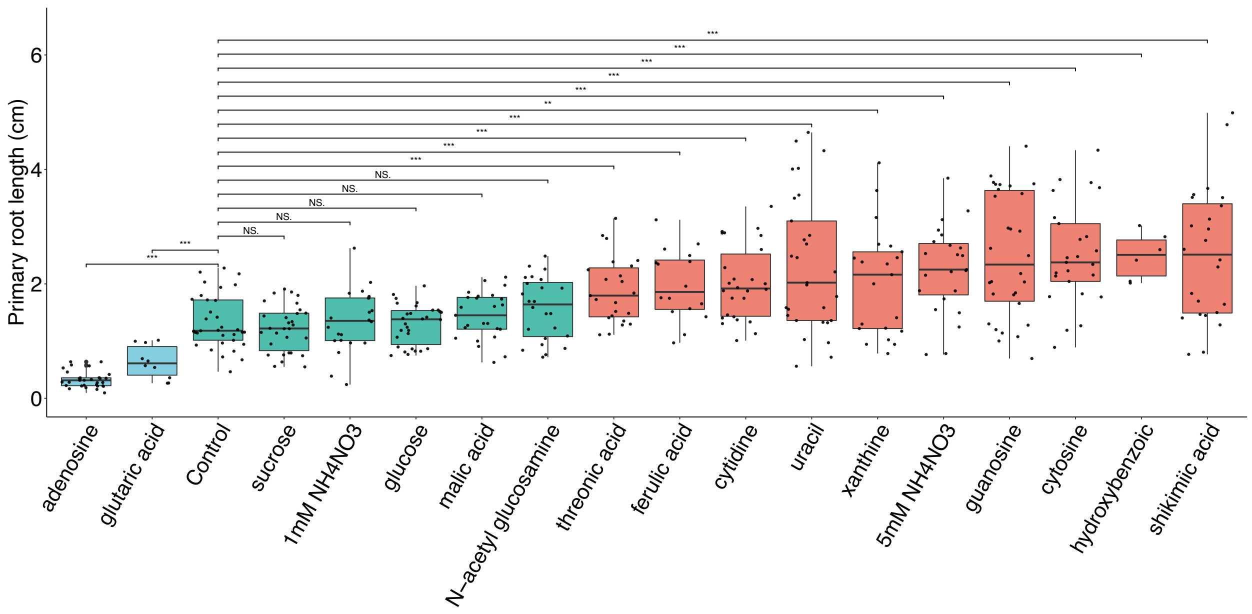


**Figure S7** ***A. thaliana*** **primary root length under addition of 0.5 mM selected metabolites and 0.5mM $NH_4NO_3$ (n=19 to 31).** Data with labels of 1mM $NH_4NO_3$ and 5mM $NH_4NO_3$ shows the root length of *A. thaliana* with high inorganic nitrogen treatment, in which plants were incubated with 1mM $NH_4NO_3$ and 5mM $NH_4NO_3$. Significance was determined via Kruskal–Wallis test followed by Dunn's test with Benjamini-Hochberg correction (NS. not significant, *$p$<0.05, **$p$<0.01, ***$p$<0.001).

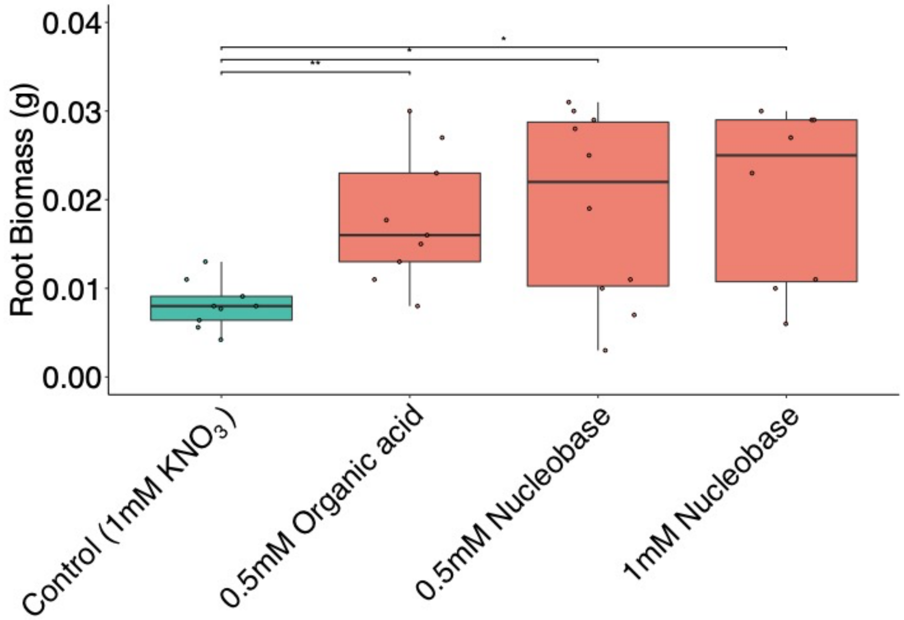


**Figure S8** ***B. distachyon*** **root biomass under addition of different classes of metabolites and inorganic nitrogen.** Control group is supplemented with 1 mM $KNO_3$ (n=9). 0.5 mM organic acid group contains 0.17 mM of threonic acid, 0.17 mM of hydroxybenzoic acid and 0.17 mM of shikimic acid (0.5 mM organic acid in total) (n=9). 0.5 mM nucleobase group contains 0.17 mM of uracil, 0.17 mM of thymine and 0.17 mM of cytosine (0.5 mM nucleobase in total) (n=10). 1 mM nucleobase group contains 0.33 mM of uracil, 0.33 mM of thymine and 0.33 mM of cytosine (1 mM nucleobase in total) (n=8). Significance was determined via Kruskal–Wallis test followed by Dunn's test with Benjamini-Hochberg correction (*$p$<0.05, **$p$<0.01, ***$p$<0.001).

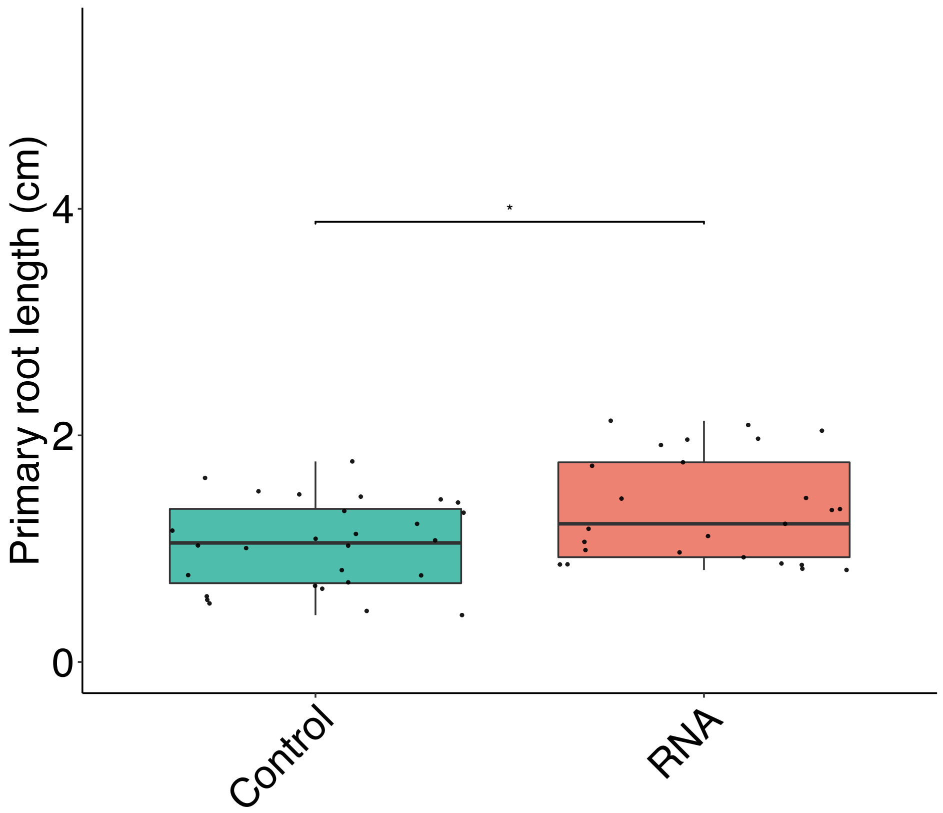


**Figure S9** ***A. thaliana*** **primary root length under addition of RNA (with N content equals to 0.5mM N) and RNase A (70 µU/mL) (n=20 to 30).** Significance was determined via Kruskal–Wallis test (*$p$<0.05).
